# The mechanosensitive ion channel Piezo1 modulates the migration and immune response of microglia

**DOI:** 10.1101/2022.06.17.496581

**Authors:** Ting Zhu, Shashwati Kala, Jinghui Guo, Yong Wu, Hui Chen, Jiejun Zhu, Kin Fung Wong, Chi Pong Cheung, Xiaohui Huang, Xinyi Zhao, Ting Lei, Minyi Yang, Lei Sun

## Abstract

Microglia are the brain’s resident immune cells, performing surveillance to promote homeostasis and healthy functioning. While chemical signaling in microglia is well-studied, the mechanical cues regulating their function are less well-understood. Here, we investigate the role of the mechanosensitive ion channel Piezo1 in microglia migration, pro-inflammatory cytokine production and stiffness sensing. We demonstrated the functional expression of Piezo1 in microglia and identified genes whose expression was affected by conditional Piezo1 knockout in transgenic mice. Functional assays revealed that Piezo1-deficiency in microglia enhanced migration towards amyloid β-protein, and decreased levels of pro-inflammatory cytokines produced upon stimulation by lipopolysaccharide, both *in vitro* and *in vivo*. The phenomenon could be mimicked or reversed using a Piezo1-specific agonist or antagonist. Finally, we also showed that Piezo1 mediated the effect of substrate stiffness-induced migration and cytokine expression. Altogether, we show that Piezo1 is an important molecular mediator for microglia, its activation modulating their migration and immune responses.

## Introduction

As the resident innate immune cells of the central nervous system (CNS), microglia play pivotal roles in maintaining brain homeostasis, pathogen defense and repairing injuries (Matcovitch-Natan, Winter et al. 2016). Microglia can recognize pathogens via pathogen-associated molecular patterns (PAMPs) on their surface (Kigerl, de Rivero Vaccari et al. 2014) and migrate to the sites of damage along a chemo-attractant gradient (Fan, Xie et al. 2017). They are able to secrete chemicals, e.g., inflammatory cytokines (Aloisi 2001) and neurotrophic factors (Nakajima, Tohyama et al. 2007) in order to protect the brain. In addition to responding to biochemical stimuli, physical cues including tissue stiffness and mechanical stimulation are also thought to contribute to microglial activity (Moshayedi, Ng et al. 2014, Bollmann, Koser et al. 2015, Ayata and Schaefer 2020). Previous studies have shown that stiff surgical implants in the CNS led to a localized foreign body reaction (FBR) by enhancing inflammatory activation of glial cells (Moshayedi, Ng et al. 2014). Putting aside the artificial manipulation of CNS microenvironmental stiffness, brain tissue itself is mechanically heterogeneous (Elkin, Azeloglu et al. 2007), and these mechanical properties may alter with age (Segel, Neumann et al. 2019) or pathological conditions (Murphy, Huston et al. 2011, Riek, Millward et al. 2012, Schregel, Wuerfel et al. 2012, Streitberger, Sack et al. 2012). Thus, microglia are exposed to varying mechanical conditions through their life cycle, but little is known about molecular mechanism how these stimuli may modulate microglial activity.

A growing body of evidence implicates the roles of the cytoskeleton, cell adhesion and Rho signaling in the process of mechanosensation and mechanotransduction (Ohashi, Fujiwara et al. 2017). However, located in lipid bilayers of cell membranes, mechanically gated ion channels serve as primary mediators of cellular mechanosensation, and merit deeper examination as well (Lim, Jang et al. 2018). Upon the increase of membrane tension, mechanically-activated ion channels undergo conformational changes and allow the flow of ions across the membrane, sensing and transducing external physical stimuli into electrochemical activity that further influences cell signaling and behavior (Mobasheri, Carter et al. 2002, Wang and Thampatty 2006). Of particular note is Piezo1, a mechanosensitive cation ion channel that detects mechanical forces with high sensitivity and broad dynamic range (Syeda, Florendo et al. 2016). Piezo1 has been reported to be expressed at high levels in endothelial cells and microglia of the brain according to previous RNA sequencing results (Zhang, Chen et al. 2014). Substantial recent research reveals that Piezo1 transduces mechanical cues to cells (Coste, Mathur et al. 2010) and is involved in many physiological and pathological process (Pathak, Nourse et al. 2014, Cahalan, Lukacs et al. 2015, Chen, Wanggou et al. 2018, Nonomura, Lukacs et al. 2018, Romac, Shahid et al. 2018, Solis, Bielecki et al. 2019). Therefore, Piezo1 is a potential mechanosensor and may be involved in coordinating the functions of microglia. In light of this evidence, we hypothesized that Piezo1 may play important roles in the functioning of microglia.

In the present study, microglia from mouse brains, *in vivo* and *ex vivo*, and a mouse microglial cell line were used as models for examining Piezo1 expression and its role in microglial functions. RT-PCR analysis showed that Piezo1 was expressed at high levels in primary microglia compared to other mechanosensitive ion channels. Gene clusters affected by the conditional knockout (KO) of Piezo1 in microglia were identified by RNA sequencing (RNAseq) and were found to include genes associated with cell motility and extracellular matrix composition. Functional analyses confirmed that Piezo1 deficiency in microglia resulted in enhanced migration ability towards Aβ_1-42_, a major component of Alzheimer’s Disease plaques. Moreover, activation of Piezo1 could suppress the migration of microglia, while blockade of Piezo1 enhanced it. We also found that microglia lacking Piezo1 showed decreased pro-inflammatory cytokines (IL-1β, IL-6 and TNF-α) production when challenged by lipopolysaccharide (LPS). Activation of Piezo1 by Yoda1, a Piezo1-specific agonist, increased production of these pro-inflammatory cytokines production by microglia when stimulated by LPS. Variation in microglial substrate stiffness was mimicked *in vitro* through modifiable gels, and we found that the expected stiffness-dependent reduction in migration and increase in pro-inflammatory cytokines production by microglia were both Piezo1-dependent. Interestingly, we also found the expression of Piezo1 was itself enhanced in microglia cultured on stiffer substrates. In summary, our study reveals that Piezo1 is a mechanosensor of stiffness in microglia, and that its activity modulates the migratory tendencies and immune response of microglia.

## Results

### Piezo1 is functionally expressed in microglial cells

We first attempted to confirm the expression of Piezo1 in primary microglia. Microglia from C57BL/6J mouse pups at postnatal day 3 (P3) were isolated by magnetic-activated cell sorting (MACS) and the purity was checked using flow cytometry (Fig. S1). The expression of various well-established mechanosensitive ion channels was then compared through RT-PCR (Ohana, Newell et al. 2009, Konno, Shirakawa et al. 2012, Shi, Du et al. 2013, Echeverry, Rodriguez et al. 2016, Cojocaru, Burada et al. 2021). Piezo1 was the channel most abundantly expressed, and its expression was significantly higher than its homolog Piezo2 (Coste, Mathur et al. 2010) (Fig. 1A). Functional analysis of Piezo1 in primary microglia cultured for 5 days was performed by patch clamping cells treated with a Piezo1-specific agonist Yoda1 (Syeda, Xu et al. 2015) and the Piezo1 blocker GsMTx4 (Bae, Sachs et al. 2011). 30 µM Yoda1 could induce strong inward current in microglia (462.7±182.1 pA), but pre-treatment with 1 µM GsMTx-4 significantly reduced the currents induced (40.4±15.36 pA) (Fig. 1B-C). Piezo1 being a cation ion channel with high permeability to Ca^2+^, calcium imaging was performed on cells loaded with a fluorescent indicator (Fura-2). We found that Yoda1 at different concentrations (2 and 10 µM) could induce robust intracellular calcium increase in primary microglia, and adding GsMTx4 significantly decreased the Ca^2+^ influx induced by 10 µM Yoda1 (Fig. 1D-F).

**Fig. 1.**
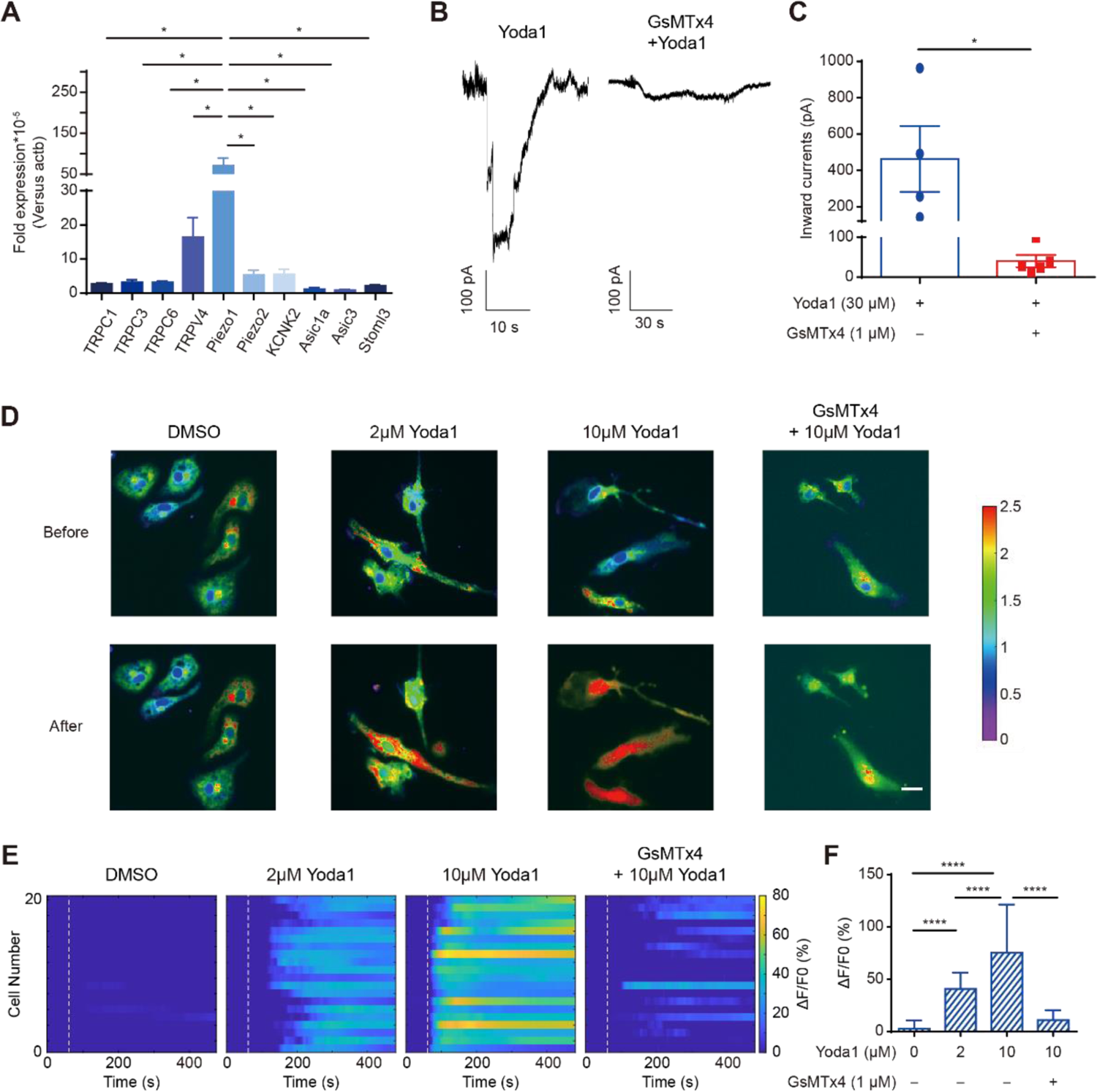
Piezo1 is functional expressed in primary microglia. **A.** The relative expression of mechanically-activated ion channel messenger RNA in primary microglia was analyzed by real-time RT-PCR using the ΔΔCT method. Bars represent mean ± SEM. N=3 independent experiments. Two-tailed unpaired t-test. *p < 0.05. **B.** Representative traces of Yoda1-induced inward currents in primary microglia recorded at −60 mV under the indicated conditions. **C.** Scatter plots of Yoda1-induced inward currents in the indicated conditions. Bars represent mean ± SEM. N=4 in Yoda1 group, n=5 in GsMTx4+Yoda1 group. Two-tailed unpaired t-test. *p < 0.05. **D.** Representative live-cell imaging of primary microglia loaded with Fura-2 at rest state (upper panel) and at the time of maximum fluorescence (lower panel). Scale bar= 20μm. **E.** Temporal raster plots of fluorescence changes in primary microglia treated with different Yoda1 and/or GsMTx4 concentrations. Dashed lines indicate when Yoda1 was added. **F.** Statistical analysis of ΔF/F0 (%) in primary microglia under different Yoda1 and/or GsMTx4 concentrations. Bars represent the mean ± SD. N= 3 independent experiments; 20-30 cells analyzed per experiment. Two-tailed Mann–Whitney U test. ****p < 0.0001.

We also tested the calcium response patterns in a commonly-used mouse microglial cell line, BV2. Yoda1 significantly increased intracellular calcium levels in BV2, and the responses were dose dependent, but GsMTx4 (1µM) inhibited the calcium increase induced by 10 µM Yoda1 (Fig. S2A-C). Thus, we concluded that Piezo1 was expressed, and that the channels were functional, in the cellular models of microglia chosen for our experiments.

### Piezo1 deficiency affects expression profiles of genes

Piezo1 has been documented to play key roles in some important physiological functions, e.g., lymphatic valve formation (Nonomura, Lukacs et al. 2018), neuronal sensing of blood pressure (Zeng, Marshall et al. 2018), and red blood cell volume homeostasis (Cahalan, Lukacs et al. 2015). Having confirmed the robust expression of Piezo1 in primary microglia, we were interested in exploring the role of Piezo1 in regulating microglial functions. Knockout or knockdown models of genes are a crucial way to study their roles in various cells. Global deletion of the Piezo1 gene is known to be lethal to embryos (Li, Hou et al. 2014); thus, we instead generated transgenic mice using a conditional knockout model. A mouse line having Piezo1 conditionally depleted from Tmem119-expressing CNS microglia was generated, of the genotype Piezo1^flox/flox^ Tmem119^CreERT2/+^ (Piezo1^ΔTmem119^) (Fig. 2A). Mice of the genotype Piezo1^flox/+^Tmem119^CreERT2/+^ (Piezo1^fl/+^) were used as a control group for comparison. Genotyping and Piezo1 knock-out validation results were performed by PCR to confirm successful deletion (Fig. S3). To evaluate the consequences of Piezo1 loss, primary microglia were isolated from brains of adult mice by MACS (Fig. 2B). Total RNA was extracted immediately after sorting and was commercially analyzed for changes in their transcriptional profile using RNAseq. The analysis revealed that 621 genes were upregulated and 293 were downregulated in Piezo1^ΔTmem119^ compared to the control (Fig. 2C-D). A gene ontology (GO) pathway analysis was performed, and the top 20 significantly enriched terms were compiled. The analysis revealed several clusters of dysregulated genes, including genes responsible for extracellular matrix organization and regulating cellular motility and migration (Fig. 2E). We thus found evidence that Piezo1 is involved in regulating cell motility, a key aspect of microglial behavior.

**Fig. 2.**
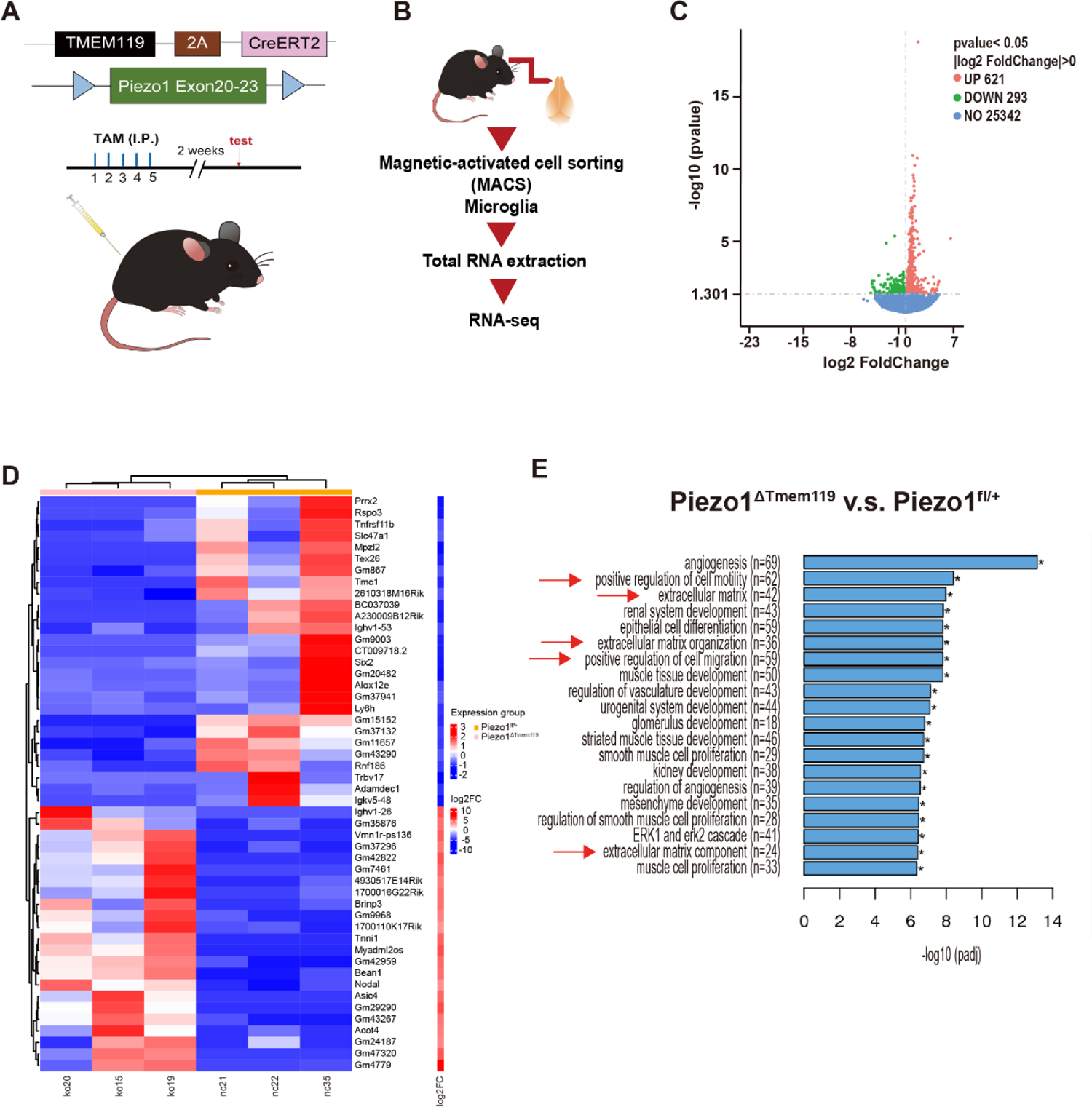
Piezo1 deficiency in microglia affects expression profiles of genes. **A.** Schematic illustration of the Piezo1 conditional knock out mice and treatment scheme with tamoxifen. **B.** Schematic illustration of RNA-seq preparation procedure. **C.** Volcano plot showing Piezo1^ΔTmem119^ vs. Piezo1^fl/+^ differentially expressed genes by RNA-seq based on primary microglia isolated from Piezo1 conditional knock out mice. **D.** Heat map showing the top 50 differentially expressed genes, Piezo1^ΔTmem119^ vs. Piezo1^fl/+^. **E.** The top 20 significantly enriched terms in the gene ontology (GO) enrichment analysis.

### Piezo1 regulates migration ability of microglia

Based on the RNAseq data, we were interested in directly investigating the impact of Piezo1 on microglial migration. We first tested whether the migration of BV2 cells could be affected by Piezo1 channel activity using an *in vitro* transwell assay. 10% FBS or 0.5 µM Aβ_1-42_, the main component of plaques in Alzheimer’s Disease, were used as chemo-attractants for BV2 cells seeded in the upper chamber of a membrane insert. Yoda1 and/or GsMTx-4 were added to the BV2 cells to modify the functioning of Piezo1. We found that the addition of Yoda1 (23.33 ± 1.202 cells towards 10% FBS, 25.67 ± 0.2404 cells towards 0.5 µM Aβ_1-42_) significantly suppressed the migratory tendency of BV2 cells compared to an untreated control (168.0 ± 8.544 cells towards 10% FBS, 51.73 ± 0.7688 cells towards 0.5 µM Aβ_1-42_), whereas adding GsMTx-4 (240.3 ± 13.35 cells towards 10% FBS, 67.93 ± 4.476 cells towards 0.5 µM Aβ_1-42_) enhanced the cells’ movement (Fig. 3A-B). The observed migration patterns were similar with both attractants used. Thus, we found that the activation or blocking of the Piezo1 channel could indeed affect the ability of BV2 cells to migrate.

**Fig 3.**
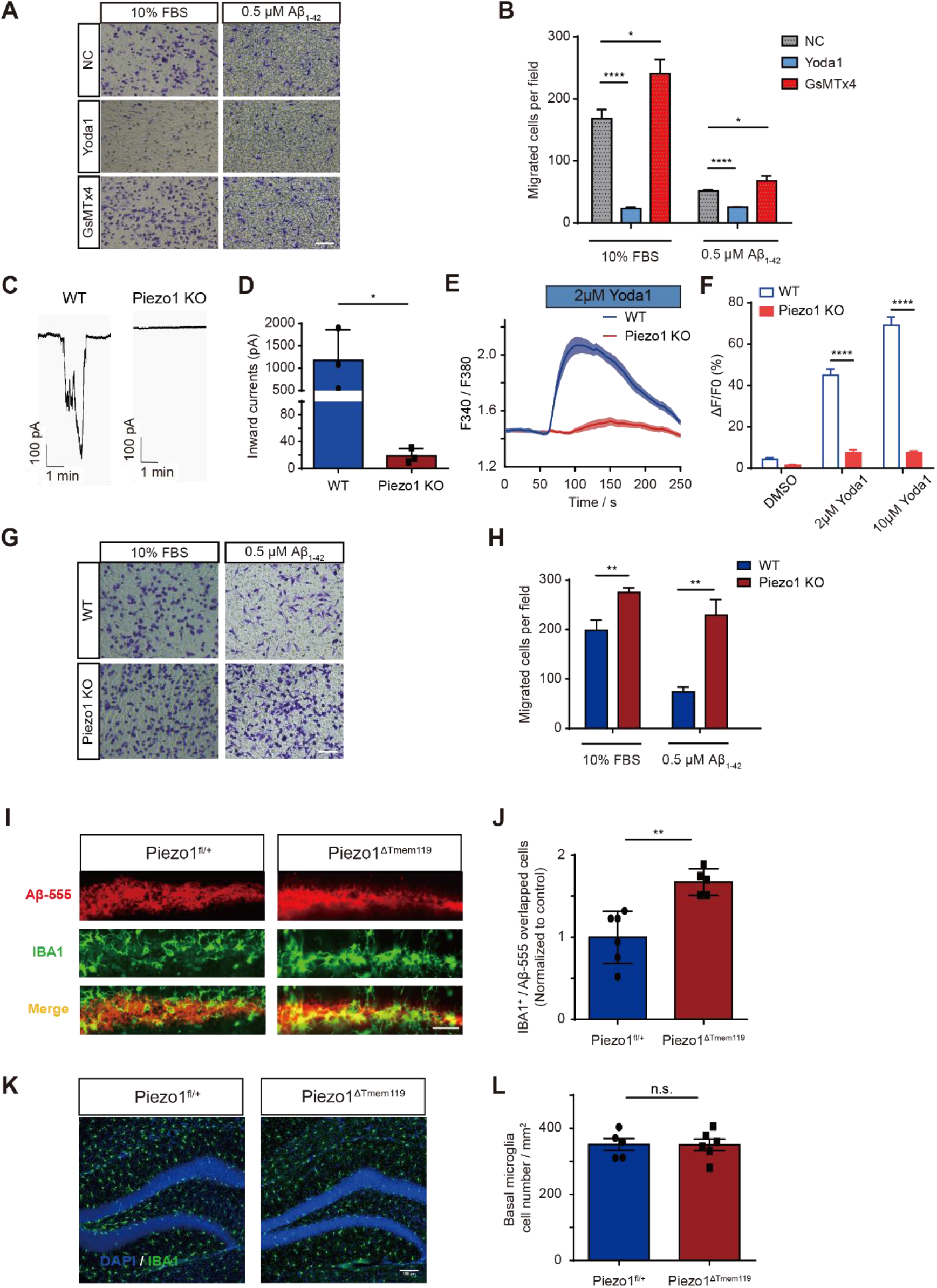
Piezo1 regulates migration ability of microglia both *in vivo* and *in vitro*. **A.** Representative images of migrated BV2 cells stained by crystal violet treated by Yoda1 (10μM) or GsMTx4 (1μM), attracted by 10% FBS or 0.5 μM Aβ_1-42_ respectively. Scale bar = 100um. **B.** Quantification of migrated cell numbers treated by Yoda1(10μM) or GsMTx4 (1μM) attracted by 10% FBS or 0.5 μM Aβ_1-42_ respectively. Bars represent mean ± SD. Repeated for 3 times. Two-tailed unpaired t test. *p < 0.05, ****p < 0.0001. **C.** Representative traces of 30μM Yoda1 induced inward currents in BV2-WT and BV2-Piezo1 KO cells recorded at −60 mV in the indicated conditions. **D.** Quantification of 30 μM Yoda1-induced inward currents in BV2 WT or BV2-Piezo1 KO cells. Bars represent mean ± SD. N=3 cells for each group. Two-tailed unpaired t test. *p < 0.05. **E.** Representative time course of calcium fluorescence in BV2-WT and BV2 - Piezo1 KO cells treated with 2μM Yoda1. **F.** Statistical analysis of ΔF/F0 (%) in BV2 WT and BV2-Piezo1 KO cells under different Yoda1 concentrations. Bars represent mean ± SEM. N=3 experiments; 30–40 cells were analyzed for each experiment. Two-tailed Mann–Whitney U test. ****p < 0.0001. **G.** Representative images of migrated BV2 WT or BV2-Piezo1 KO cells stained by crystal violet, attracted by FBS or Aβ_1-42_ respectively. Scale bar=100um. **H.** Quantification of migrated BV2 WT or BV2-Piezo1 KO cell numbers attracted by FBS or Aβ_1-42_ respectively. Bars represent mean ± SD. Repeated for 3 times. Two-tailed unpaired t test. **p < 0.01. **I.** Representative images of Aβ_1-42_ injected regions in the dentate gyrus (DG) region of Piezo1^fl/+^ and Piezo1^ΔTmem119^ mouse groups respectively. Scale bar = 50μm. **J.** Quantified average values for total Aβ-555/IBA1 positive (IBA1+) cells, normalized to control group (n=6 in Piezo1^fl/+^ group, n=5 in Piezo1^ΔTmem119^ group). Bars represent mean ± SD. Two-tailed unpaired t test. **p < 0.01. **K.** Representative images of IBA1 positive cells in DG area of Piezo1^fl/+^ and Piezo1^ΔTmem119^ brain slice respectively (blue: DAPI, green: IBA1). Scale bar= 100μm. **L.** Quantification of microglia numbers per mm2 around DG area in Piezo1fl/+ and Piezo1ΔTmem119 mice respectively (n=5 in Piezo1fl/+ group, n=6 in Piezo1ΔTmem119 group). Bars represent mean ± SEM. Two-tailed unpaired t test. n.s., no significance.

We next constructed a BV2-Piezo1 Knock-out (KO) stable cell line using CRISPR-Cas9 technology. We used a patch clamp setup to confirm that BV2-Piezo1 KO cells were insensitive to Yoda1 stimulation compared to WT cells (Fig. 3C). Inward current induced by 30 µM Yoda1 in BV2-WT cells was 1176 ± 394.8 pA while it was much lower (18.67 ± 6.333 pA) in BV2-Piezo1 KO cells (Fig. 3D). The resting membrane potential of BV2 cells with Piezo1 KO was not obviously different from wild type cells (Fig. S4) indicating that Piezo1 KO did not itself alter the cells’ general electrical activity and health. Calcium imaging also showed that BV2-Piezo1 KO cells were insensitive to Yoda1 stimulation (2µM and 10µM) (Fig. 3E-F). Piezo1 depletion was also found to significantly enhance the cells’ migration towards both FBS and Aβ_1-42_ compared to WT (Fig. 3G-H). The cell migration process requires close control and coordination of actin dynamics and actin signaling dysfunction has been known to lead to aberrant cell migration (Schaks, Giannone et al. 2019). Hence, we also examined the effect of Piezo1 KO on the polymerization of actin. Yoda1 was seen to significantly enhance actin polymerization in WT cells, but this effect was abolished in KO cells (Fig. S5A-B). Interestingly, Piezo1 KO was found to reduce basal actin polymerization levels. Thus, knocking out Piezo1 in BV2 cells was found to significantly affect the cells’ electrical responses, calcium dynamics and tendency to migrate.

The role of Piezo1 in mediating the migration ability of microglia was further examined by a simple chemoattraction assay *in vivo*. The oligomer Aβ_1-42_ HiLyte™ Fluor 555 was injected into the dentate gyrus (DG) of the hippocampus as an attractant. Counting IBA1-positive cells inside the Aβ_1-42_ −555 signal regions, many more microglia were found to have migrated to the DG in Piezo1^ΔTMEM119^ mice than control Piezo1^fl/+^ mice (Fig. 3I-J) by. We also counted microglia in mice of both genotypes without any injection of Aβ_1-42_, and found that conditional Piezo1 KO did not affect the microglia numbers in their basal state (Fig. 3K-L). Thus, we found that Piezo1 expression and activation regulated of the ability of microglia to migrate towards attractant stimuli *in vitro* and *in vivo*.

### Piezo1 regulates the immune response of microglia

Since microglia act as key players in the primary immune response of the CNS, we also wanted to investigate whether Piezo1 was involved in the microglial immune response. For this, we used BV2-Piezo1 KO and WT cell lines, stimulated with Yoda1 and/or LPS for 24 hours, a toxin used to trigger an inflammatory response. The expression levels of some important pro-inflammatory cytokines were then monitored by RT-PCR. In BV2-WT cells stimulated with LPS, we found that Yoda1 increased IL-1β, IL-6 and TNF-α mRNA expression levels (Fig. 4A-C). We also found that BV2-Piezo1 KO cells expressed significantly lower levels of these cytokines even when stimulated with LPS (Fig. 4D-F). Western blotting for IL-1β further backed up the finding that loss of Piezo1 expression significantly reduced the ability of microglia to response to LPS stimulation by producing cytokines (Fig. 4G-H). We further tested the immune response mediation by Piezo1 *in vivo* using our transgenic mouse model. The cortex tissues of mice were evaluated for cytokine expression levels after intraperitoneal LPS injection. Mice with the conditional Piezo1 KO in microglia showed lower expression of these cytokines when challenged by LPS, but not in the basal state (Fig. 4I-K). Piezo1 was hence seen to play crucial roles in the LPS-induced neuroinflammatory response of microglia.

**Fig 4.**
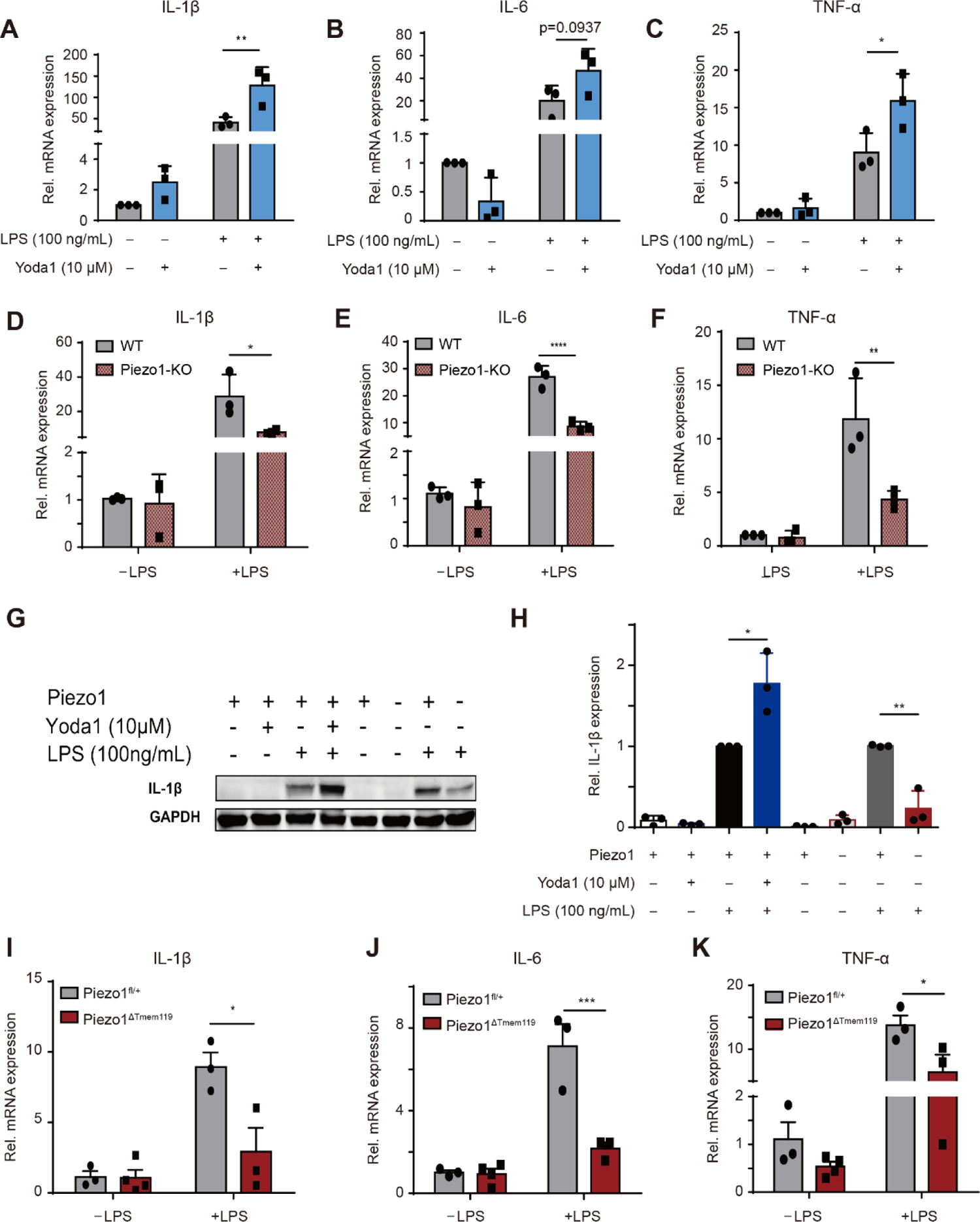
Piezo1 regulates production of pro-inflammatory cytokines by microglia when challenged by LPS both *in vivo* and *in vitro*. **A-C.** Relative mRNA expressions of IL-1β, IL-6 and TNF-α in BV2-WT cells, exposed to Yoda1 (10µM) or not, in basal state or stimulated by LPS (100ng/mL), normalized to BV2 basal control state. Bars represent mean ± SD. N=3 independent experiments. *p < 0.05, **p < 0.01. **D-F.** Relative mRNA expressions of IL-1β, IL-6 and TNF-α of BV2-WT and BV2-Piezo1 KO cells in basal state or stimulated by LPS (100ng/mL), normalized to BV2-WT basal state. Bars represent mean ± SD. N=3 independent experiments. *p < 0.05, **p < 0.01, ****p < 0.0001. **A. G.** Representative Western blot images for IL-1 β and GAPDH in indicated conditions. **B. H.** Quantification of IL-1 β protein levels normalized to GAPDH compared with BV2-WT treated with LPS group. Bars represent mean ± SD. N=3 independent experiments. Two-tailed unpaired T test. *p < 0.05, **p < 0.01. **I-K.** Relative mRNA expressions of IL-1β, IL-6 and TNF-α in cortex of Piezo1^fl/+^ and Piezo1^ΔTMEM119^ mice in basal state or after administration of LPS (n=3 in Piezo1^fl/+^ - LPS group, n=4 in Piezo1^ΔTmem119^-LPS group, n=3 in Piezo1^fl/+^ +LPS group, n=3 in Piezo1^ΔTmem119^ +LPS group). Bars represent mean ± SEM. For panel a-f and i-k, One-way ANOVA with post-hoc Tukey test. *p < 0.05, ***p < 0.001.

### Stiffness-mediated microglial migration and activation are mediated by Piezo1

Given that Piezo1 is involved in mechanosensation in many cell types and physiological functions (Li, Hou et al. 2014, Pathak, Nourse et al. 2014, Cahalan, Lukacs et al. 2015, Gudipaty, Lindblom et al. 2017, Romac, Shahid et al. 2018, Solis, Bielecki et al. 2019), we reasoned that Piezo1 would play such a role in microglia as well. To investigate the role of Piezo1 in microglia sensing of stiffness, to regulate migration ability and cytokine production of microglia, BV2-WT and BV2-Piezo1 KO cells were plated onto soft (0.6 kPa) or stiff (35 kPa) polyacrylamide gel substrates (as previously described (Chang, Chen et al. 2017)) for 48 hours. The cells’ migration towards 10% FBS and expression of inflammatory cytokines were then compared between groups. We found that enhanced substrate stiffness reduced the migration of BV2-WT cells but not in BV2-Piezo1 KO cells (Fig. 5A-B). We also found that greater substrate stiffness could increase the expression of pro-inflammatory factors (IL-1β, IL-6 and TNF-α) in BV2-WT cells, but did not in IL-1β and TNF-α expression in BV2-Piezo1 KO cells (Fig. 5C-E), indicating Piezo1 is involved in stiffness-induced microglial migration and activation process.

**Fig 5.**
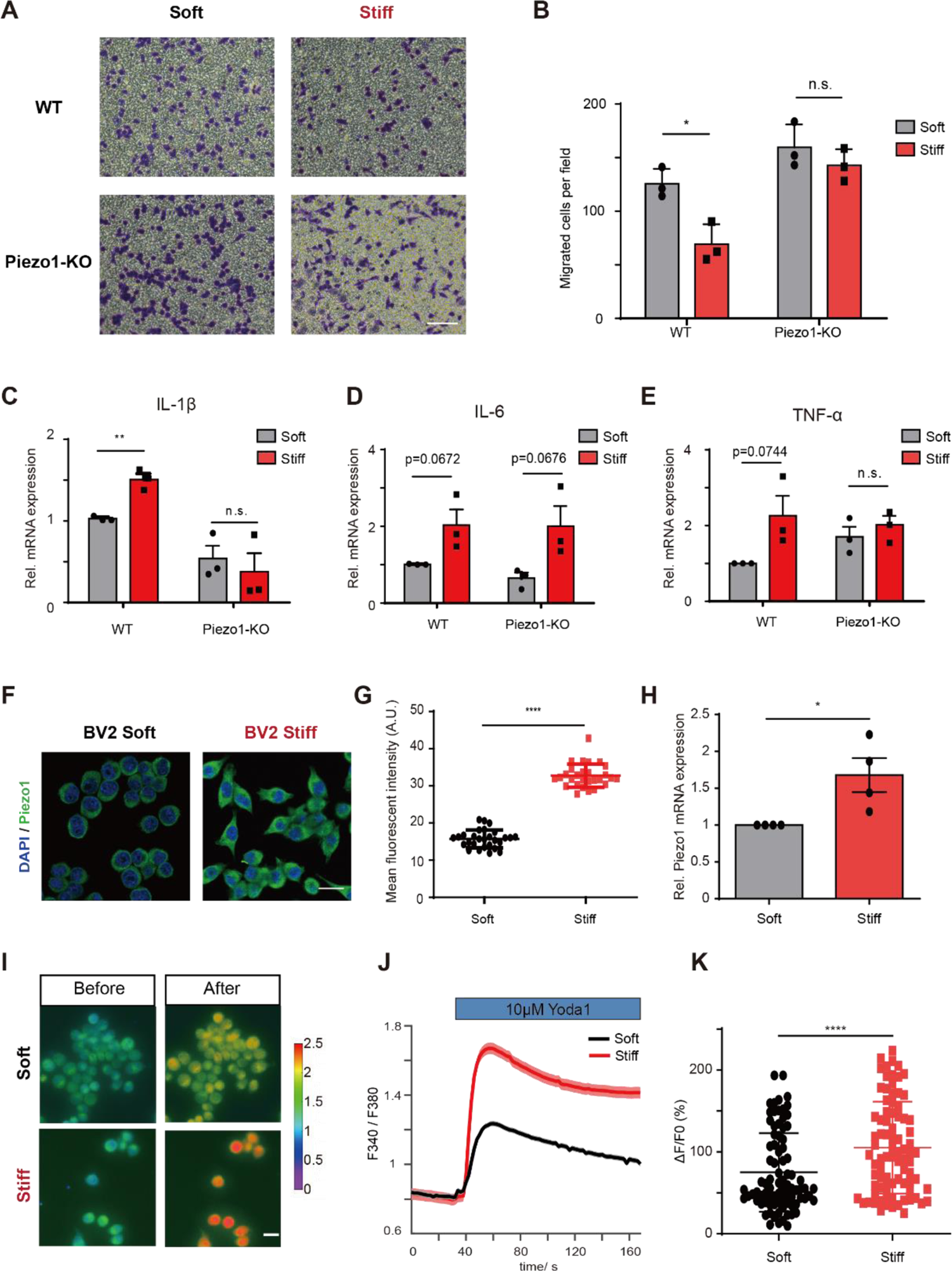
Stiffness-dependent Piezo1 expression/activity modulates microglia pro-inflammatory cytokines production. **A.** Representative images of migrated BV2 WT or Piezo1-KO cells pre-cultured for 48h in PA gel of varying stiffness (Soft: 0.6KPa, Stiff: 35KPa) stained by crystal violet. Scale bar = 100um. **B.** Quantification of migrated cell numbers of BV2 WT or Piezo1-KO cells pre-cultured for 48h in PA gel of different stiffness (Soft: 0.6KPa, Stiff: 35KPa). Bars represent mean ± SD. N=3 independent experiments. Two-tailed unpaired t test. n.s., no significance, *p < 0.05. **C-E.** Relative IL-1β, IL-6 and TNF-α mRNA expression of BV2-WT and BV2-Piezo1 KO seeded on surfaces of indicated stiffness (Soft: 0.6KPa, Stiff: 35KPa). Bars represent mean ± SEM. N=3 independent experiments. Two-tailed unpaired t test. n.s., no significance; **p < 0.01. **F.** Immunofluorescence confocal images of Piezo1 (green) in BV2 cell lines seeding on different stiffness of PA gel (Soft: 0.6KPa, Stiff: 35KPa). Scale bar= 20μm. **G.** Quantification of Piezo1 mean fluorescent intensity (MFI) of BV2 cell lines seeding on different stiffness of PA gel. Bars represent mean ± SD. Repeated for 3 times, around 30 cells were analyzed for each experiment. Two-tailed unpaired t test. ****p < 0.0001. **H.** Relative Piezo1 mRNA levels of BV2 cell lines seeding on different stiffness of PA gel. Bars represent mean ± SEM. N=4 independent experiments. Two-tailed unpaired t test. *p < 0.05. **I.** Representative live-cell imaging of BV2 cells loaded with Fura2 calcium indicator treated by Yoda1 (10μM) at time 0 (left panel) and at the time of maximum fluorescence (right panel) seeding on soft and stiff substrate respectively. **J.** Representative time course of calcium fluorescence in BV2 cells seeding on soft (black curve) and stiff substrates (red curve) treated with Yoda1 (10μM). **K.** Statistical analysis of ΔF/F0 (%) of BV2 cells seeding on soft (black) and stiff substrates (red) induced by Yoda1 (10μM). Bars represent mean ± SD. N=3 independent experiments, 20-30 cells were analyzed for each experiment. Two-tailed Mann–Whitney U test. ****p < 0.0001.

An unexpected aspect of the soft/stiff substrate culture was that an increase in stiffness was also found to be correlated with increased Piezo1 expression. BV2 cells on a stiffer substrate showed significantly increased Piezo1 expression after 48 hours of culture in both protein levels (Fig. 5F-G) and mRNA levels (Fig. 5H). Stiff-cultured cells also showed much greater calcium response when stimulated with 10 µM Yoda1 than did soft-cultured cells (Fig. 5I-K), which also points towards an increased expression of Piezo1. Thus, we found that Piezo1 was both a mediator for the microglial response to environmental stiffness cues, while also being upregulated in microglia in response to varying mechanical conditions.

## Discussion

In the CNS, microglia constantly encounter mechanically heterogeneous regions of varying cellular density and matrix composition. This implies the presences of a suite of cellular machinery to sense and transduce these cues into cellular signaling. In the present study, we identify a role for the mechanically-activated ion channel Piezo1 in regulating microglial functions, i.e., migration, pro-inflammatory cytokines production and stiffness sensing. We found that Piezo1 is highly expressed in primary cultured microglia. Microglia from mice with conditional Piezo1 knockout exhibited enhanced migration ability towards injected Aβ_1-42_ and reduced pro-inflammatory cytokines production after stimulation by LPS. These phenomena were also observed *in vitro* using constructed BV2-WT and BV2-Piezo1 KO stable cell lines. I*n vitro*, we found that activation and blockade of the Piezo1 channel reversely regulated the migration tendencies of microglia. Furthermore, Yoda1 could enhance pro-inflammatory cytokines production of microglia stimulated by LPS, but similarly activating Piezo1 in unstimulated microglia had little effect. We also show that substrate stiffness modulates Piezo1 expressions and stiffness regulated microglia migration and pro-inflammatory cytokines production (IL-1β, TNF-α) enhancement was affected by Piezo1 depletion. Together, our study reveals critical roles of Piezo1 in modulating microglial functions and indicates its significance in maintaining brain health.

Ca^2+^-mediated signals are a common signal transduction pathway in all living cells, including microglia (Moller 2002). Ca^2+^ activities of microglia are known to be altered in some pathological conditions, such as interrupted Ca^2+^ signal transduction following brain damage (Eichhoff, Brawek et al. 2011) or in Alzheimer’s disease (AD) patients (McLarnon, Choi et al. 2005), though the mechanism is unclear. We found stiffness-driven variation in Piezo1 expression using our *in vitro* PA gel model. A previous study using a TgF344-AD transgenic model also found that Piezo1 expression level in astrocyte increases around stiff amyloid plaques (Velasco-Estevez, Mampay et al. 2018). Given that mechanical cues are altered in the brain during progression of several pathological conditions in which microglia are involved, e.g., traumatic brain injury (Keating and Cullen 2021) or AD (Murphy, Huston et al. 2011, Murphy, Jones et al. 2016), aberrant Piezo1expression may be a contributor to microglial calcium signaling dysfunction in such conditions as well. Thus, the role of Piezo1 in microglia, regulated by changing microenvironment stiffness of the brain, merits further study using appropriate animal models.

Resting microglia are highly dynamic and can be rapidly activated to participate in pathological responses, including migrating to affected sites, releasing various inflammatory molecules, and clearing cellular debris (Kreutzberg 1996, Koizumi, Shigemoto-Mogami et al. 2007) to maintain brain health. Therefore, migration is a critical first step for microglia to function after injury or inflammatory stimuli. Our findings identify a potentially crucial role of Piezo1 in modulating microglial migration. Numerous studies have shown that microglia tend to migrate towards chemo-attractant stimuli (Fan, Xie et al. 2017). Based on our observation that Piezo1 serves as a mechanosensor and plays a part in regulating microglial migration, mechanical cues exerted on the cells during ageing (Segel, Neumann et al. 2019) and some pathological conditions (Murphy, Huston et al. 2011, Riek, Millward et al. 2012, Schregel, Wuerfel et al. 2012, Streitberger, Sack et al. 2012) thus also merit exploration.

Piezo1 also played an important role in microglial production of pro-inflammatory cytokines (IL-1β, IL-6, TNF-α), which are involved in the pathogenesis of many neurodegenerative diseases (Smith, Das et al. 2012). Enhanced stiffness, mediated by Piezo1, was also associated with increased pro-inflammatory cytokine production in microglia. These cytokines play potentially contradictory roles in the maintenance of brain homeostasis. On one hand, release of these factors is crucial to prevent further damage to CNS tissue when, say, under attack by pathogens. However, on the other hand, the same cytokines may also be toxic to neurons and other glial cells, and contribute to the progression of neurodegenerative disorders (Smith, Das et al. 2012, Ogunmokun, Dewanjee et al. 2021, Tansey, Wallings et al. 2022). Interestingly, substrate stiffness could also regulate Piezo1 expression in microglia and induce following microglial function changes (Fig 5). This phenomenon suggests Piezo1 could be a key molecule mediating between microenvironment stiffness and microglial function. Given the mechanical cues generated during the progression of brain diseases, Piezo1 could be an important subject of study as a participant in signaling pathways and as a druggable target. Several studies have already described the roles of mechanosensitive ion channels in regulating brain diseases, i.e., TRPC1 (Wu, Ryskamp et al. 2018), TRPC3, TRPC6 (Wang, Lu et al. 2015, Chen, Lu et al. 2017), TRPV4 (Wang, Zhou et al. 2019). Given that Piezo1 appears to mediate crucial microglial functions, further study of its role in the brain could improve our understanding of the progression of brain diseases and the function of microglia in health and disease.

## Materials and methods

### Transgenic mouse line generation and genotyping

All animal procedures were approved by the Animal Subjects Ethics Sub-Committee (ASESC) of the Hong Kong Polytechnic University, and were performed in compliance with the guidelines of the Department of Health - Animals (Control of Experiments) of the Hong Kong S.A.R. government. Generation of Piezo1^ΔTMEM119^ mice was accomplished through breeding Piezo1^flox/flox^ (Jackson Laboratories Stock No. 029213) and Tmem119-2A-Cre^ERT2^ (Jackson Laboratories Stock No: 031820) mice together to generate progeny that were heterozygous for both genes. The generated heterozygotes were then bred with Piezo1^flox/flox^ mice to generate Piezo1^flox/flox^Tmem119^Cre/+^ (Piezo1^ΔTmem119^) and Piezo1^flox/+^Tmem119^Cre/+^ (Piezo1^fl/+^) mice. Mice were housed in a 12-h light/12-h dark cycle, with temperatures between 20 °C and 24 °C and 40–70% humidity with food and water available *ad libitum*. Tamoxifen (Sigma-Aldrich, T5648) was dissolved in corn oil containing 5% ethanol at 20 mg/ml under vortex at times for several hours in the dark. At time of administration, the mice weighed 19–26 g and intraperitoneal (i.p.) injection with 100μl of the 20 mg/ml tamoxifen solution for 5 consecutive days. After administration of tamoxifen for at least 2 weeks, mice were used for different *in vivo* assays. Mice tail samples were genotyped using the following primers suggested by Jackson Laboratories: Piezo1-28247-forward: GCC TAG ATT CAC CTG GCT TC; Piezo1-28248-reverse: GCT CTT AAC CAT TGA GCC ATC T; TMEM119-16504: ATC GCA TTC CTT GCAAAA GT; TMEM119-42648: CAG TAT GTG GGG TCA CTG AAG A; TMEM119-42649: ACT TGG GGA GAT GTT TCC TG using Phire Tissue Direct PCR Master Mix (F170S, Thermo Scientific) with the following cycling conditions: Initial denaturation 98°C for 5 min, followed by 10 cycles of 98°C for 5 s, 65°C for 5 s, −0.5 C per cycle decrease, 68°C for 20 s, followed by 30 cycles of 98°C for 5 s, 60°C for 5 s, 72°C for 20 s, followed by a final hold of 72°C for 1 min. Reactions were separated on 2% agarose gels yielding the following band sizes: Cre +/-: 280bp and 378bp, Cre -/-: 378bp, Piezo1 fl/fl: 380bp, Piezo1 fl/+: 380bp and 188bp. Successful knock-out Piezo1 were validated by examining DNA from mice brains using the following primers: Piezo1-28247-forward: GCC TAG ATT CAC CTG GCT TC; Piezo1 KO-reverse: AGG TTG CAG GGT GGC ATG GCT CTT TTT. An obvious Piezo1 KO band was found in 230bp but none in control mice group. Brain samples were also collected for testing Piezo1 mRNA levels by qPCR using the following primers: Piezo1-192-forward: CTCTGGCCTAACTACTGTCT; Piezo1-192-reverse: GATAAGGTTGGTGGAGTTGG.

### Cell culture

The BV2 cell line (RRID: CVCL_0182) was a gift from Dr. Bo Peng’s group. Cells were cultured in DMEM (10566016, Gibco), 10% fetal bovine serum (A3840402, Gibco), 1% Penicillin-Streptomycin (15140122, Gibco) at 37℃ inside a standard cell culture incubator containing 5% CO_2_.

### Primary microglia isolation

For primary microglia isolation, P3 pups of C57BL/6J mice were sacrificed, followed by dissociation and isolation using the CD11b isolation kit according to the manufacturer’s protocol (130-092-628 and 130-126-725, Miltenyi Biotec). Primary microglia purity was examined by flow cytometry after staining by Cd11b-FITC (130-113-243, Miltenyi Biotec) and Cd45-APC (130-110-660, Miltenyi Biotec). Dishes were pre-coated with 50μg/μl Poly-L-Lysine and cells were then plated on the dishes at a density of 10^5^ cells/Ml and cultured in DMEM/F12 (10565018, Gibco), 10% fetal bovine serum (A3840402, Gibco), 1% Penicillin-Streptomycin (15140122, Gibco) at 37℃ inside a standard cell culture incubator containing 5% CO_2_.

### CRISPR knock-out (KO) Piezo1 cell cloning

To generate Piezo1 knock-out (KO) cell lines, single guide RNAs (sgRNAs) ACGCTTCAATGCTCTCTCGC targeting the second exon of mouse piezo1 as reference (Del Marmol, Touhara et al. 2018) was inserted into MLM3636 vector (Addgene Plasmid #43860). SgRNA inserted MLM3636 and Cas9 expression plasmid pX459 v2.0 (Addgene Plasmid #62988) were transfected into BV2 cell line with Lipofectamine^TM^ 3000 transfection reagent (L3000015, Thermo Fisher Scientific). Transfected BV2 cells were selected with 1ug/mL puromycin (A1113803, Life Technologies). After selection, live cells were trypsinized, diluted as 5 cell/mL and then seeded on 96-well plate at an average density of 0.5cell/well for growing to obtain single clones. The successful knock-out clone was verified using Sanger sequencing, patch clamp and calcium imaging.

### Intracellular calcium imaging

Cells were plated on confocal dishes (200350, SPL Life Sciences). Prior to recording, cells were incubated with 2 μM Fura-2 (F1221, Invitrogen) at 37℃ for 30 min dispersed in Margo solution with 0.2‰ w/v Pluronic F-127. Cells were then washed twice and incubated for another 15 min. GsMTx-4 (ab141871, abcam) were added in this step as needed. Experiments were performed at room temperature using Nikon Eclipse Ti2-E microscope. Yoda1 (SML1558, Sigma Aldrich) or DMSO (D12345, Invitrogen) was added during the imaging process. Intracellular calcium concentration was indicated as the ratio of fura-2 emission (510 nm) intensities for 340 and 380 nm excitation.

### Patch clamp

Cells were seeded in culture dishes before patch clamp recording. Borosilicate glass-made patch pipettes (Vitrex, Modulohm A/S, Herlev, Denmark), were pulled with micropipette puller (P-97, Sutter Instrument Co., USA) to a resistance of 2 to 5 MΩ before being filled with KCl pipette solution (in mM): KCl 138, NaCl 10, MgCl_2_ 1 and HEPES 10 with D-manitol compensated for osm290. Inward current was recorded with a data acquisition system (DigiData 1322A, Axon Instruments) and an amplifier (Axopatch-200B, Axon Instruments, Foster City, CA, USA). The command voltages were controlled by a computer equipped with pClamp Version 9 software. For cell resting membrane potential (RMP) test, cells were placed on bath solution (in mM): NaCl 130, MgCl_2_ 2, KCl 4.5, Glucose 10, HEPES 20, and CaCl_2_ 2, pH 7.4. When the whole-cell Giga seal was formed and the capacitance of cell was measured, the RMP of cells was recorded when the injection current was set at zero. Afterwards, the inward currents induced by Yoda1 was recorded by voltage clamp gap free recording mode.

### RNA isolation and qPCR

RNA was extracted using the MiniBEST Universal RNA Extraction Kit (Cat No. 9767, TaKaRa). cDNA syntheses were generated using PrimeScript RT Master Mix (RR036A, TaKaRa). SYBR® Premix Ex Taq™ II (Tli RNaseH Plus) (RR820A, TaKaRa) were performed with the CFX96 Touch Deep Well Real-Time PCR system under standard conditions. The ΔΔCt method was used to calculate gene expression. Primer sequences are provided in the following table.

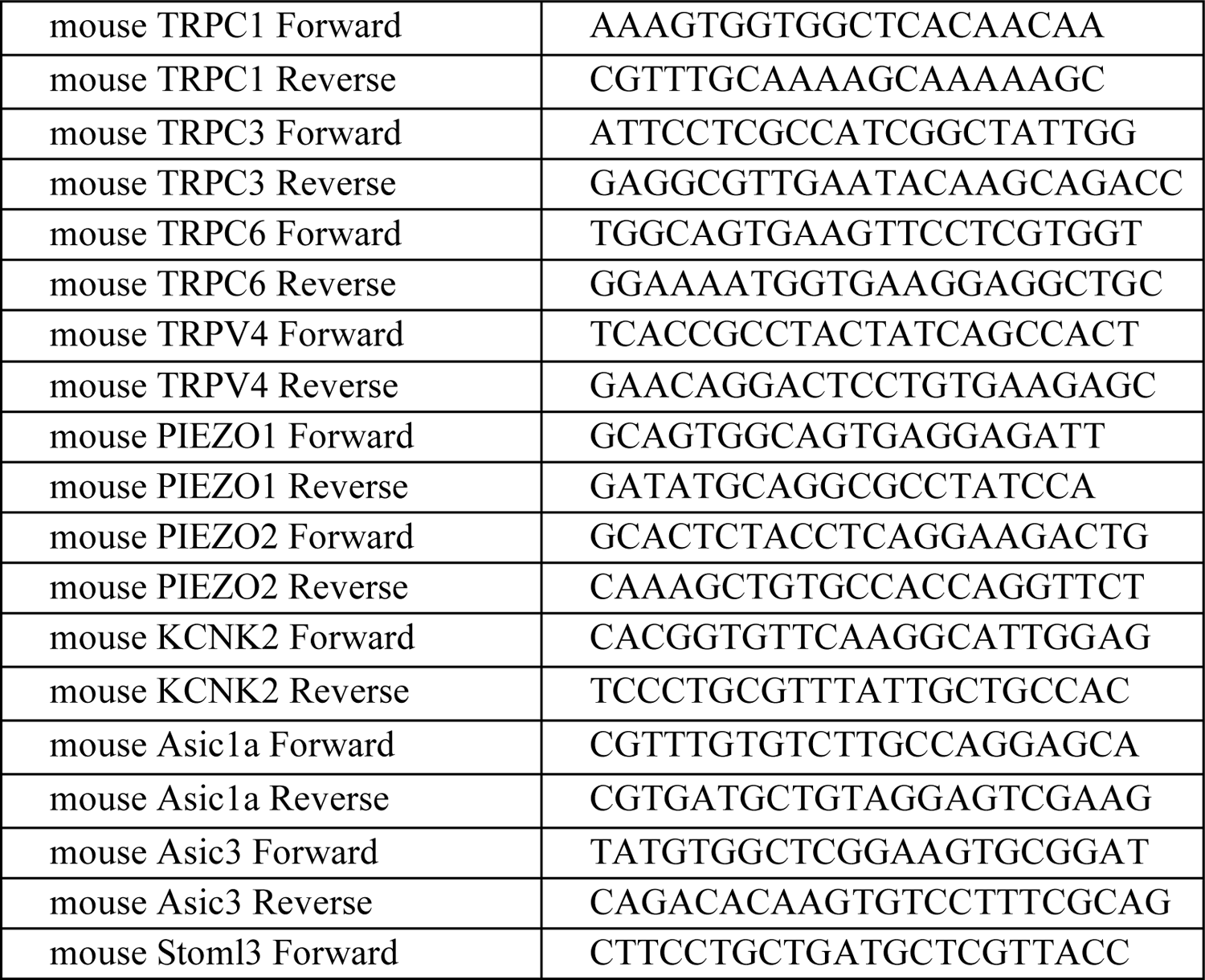
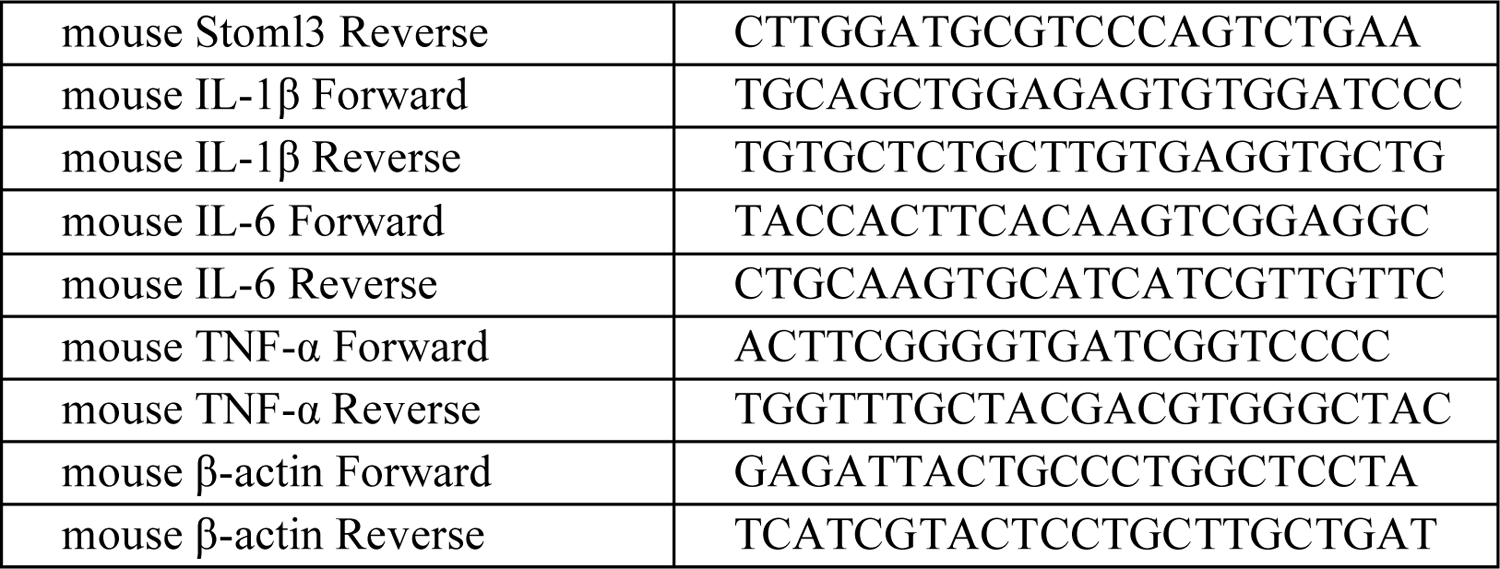

### RNA sequencing and data analysis

RNA sequencing and informatics analysis were performed by Novogene Company (Beijing, China). RNA integrity was assessed with the Agilent 2100 Bioanalyzer (Agilent Technologies). Libraries were barcoded, quantified using the NEBNext Ultra RNA Library Prep Kit for Illumina (Cat No. 7530, New England Biolabs (NEB)). Quantified libraries were pooled and sequenced on Illumina platforms, according to effective library concentration and data amount. The clustering of the index-coded samples was performed according to the manufacturer’s instructions. After cluster generation, the library preparations were sequenced on an Illumina platform and paired-end reads were generated. Raw data (raw reads) of fastq format were first processed through in-house perl scripts. Clean data (clean reads) were obtained by removing reads containing adapter, reads containing ploy-N and low quality reads from raw data. At the same time, Q20, Q30 and GC content the clean data were calculated. All the downstream analyses were based on the clean data with high quality. Reference genome and gene model annotation files were downloaded from genome website directly. Index of the reference genome was built using Hisat2 v2.0.5 and paired-end clean reads were aligned to the reference genome using Hisat2 v2.0.5. Feature Counts v1.5.0-p3 was used to count the reads numbers mapped to each gene. Differential expression analysis of two groups (3 biological replicates per condition) was performed using the DESeq2 R package. Genes with an adjusted P value <=0.05 found by DESeq2 were assigned as differentially expressed. Gene Ontology (GO) enrichment analysis of differentially expressed genes was implemented by the cluster Profiler R package, in which gene length bias was corrected. GO terms with corrected P value less than 0.05 were considered significantly enriched by differential expressed genes.

### Cell migration transwell assays

BV2 cells and primary microglia migration towards 10% FBS were tested in transwell assays (COSTAR 24-well plate with inserts, 8-μm pore, #3422, Corning, NY). For the transwell assay, 600 μl DMEM together with the attractant-10% FBS or 0.5μM Aβ_1-42_ were added below the membrane inserts. Drugs-1‰ DMSO, 10 μM Yoda1, 1 μM GsMTx4 - were respectively added to 5 × 10^4^ cells in 100 μl media right before adding the cells to the top wells. Migration was assayed for 20 h at 37℃ and 5% CO_2_. Subsequentlly, inserts were removed and washed three times with PBS. BV2 cells were fixed and stained with 0.5% crystal violet for 20min. Stationary cells were removed from the top with cotton tips. Five random fields were selected for imaging, and the migrated cells were quantified using Image J software.

### Stereotactic injection of Aβ_1-42_

Beta-Amyloid (1-42), HiLyte™ Fluor 555-labeled (Cat. AS-60480-01, ANASPEC) were dissolved in DMSO to obtain a 1 mM stock, resuspended in PBS at 100 μM, and incubated at 37 °C for 1 day to promote Aβ_1-42_ aggregation. Mice were anesthetized by i.p. injection of a mixture of ketamine (100 mg/kg), xylazine (10 mg/kg), 2 μl of Aβ_1-42_-555 was distributed into DG area of the hippocampus (AP −2 mm; ML −1.5 mm; DV 2 mm) using stereotaxic equipment (RWD Life Science Co., Ltd). After recovery from surgery, animals were returned to their home cages. Post surgery (16 h), mice were anesthetized and perfused for subsequent experiments.

For histological analyses of microglia migration towards Aβ_1-42_-555, brains were perfused with ice-cold PBS, fixed in 4% PFA and immersed in 4% PFA for another 4 hours. Brains were then sectioned in 50μM thickness by Vibrating Microtome (Leica VT1200S). Sections were washed, then permeabilized with 0.3% Triton X-100 (Sigma) in PBS with 5% Goat serum (16210072, Gibco) and blocked for 1 hour. The microglia-specific antibody IBA1 (Rabbit mAb 17198S, CST, RRID: AB_2820254) was incubated in blocking buffer overnight at 4°C followed by intensive washing and incubation with a Goat anti-rabbit Adsorbed Secondary Antibody, Alexa Fluor 488 (A-11008, Thermo Fisher Scientific, RRID: AB_143165) for 2 hours. Sections were washed again and mounted with Mounting Medium With DAPI-Aqueous, Fluoroshield (ab104139, Abcam). Data acquisition was performed using a Nikon Eclipse Ti2-E Live-cell Fluorescence Imaging System in Z-stack model.

### Western blotting

Cells were rinsed with PBS before being exposed to lysis buffer, a combination of RIPA lysis buffer (89900, Thermo Fisher Scientific) and 1% Halt™ Protease and Phosphatase Inhibitor Cocktail (78446, Thermo Fisher Scientific). Cells were scraped to obtain the adhered cells and the lysate was collected. The lysate was put on ice for another 30 min to achieve thorough lysis. Lysate was spun at 13000 rpm for 20min at 4°C and the supernatant was collected. Samples were quantified using Pierce™ Rapid Gold BCA Protein Assay Kit (A53225, Thermo Fisher Scientific) and aliquots of 50μg of protein were prepared.The proteins were denatured through the use of a loading buffer supplemented with 5% 2-mercaptoethanol at 95°C for 10 min before each sample was loaded. Gel electrophoresis resulted in the separation of proteins before being transferred onto nitrocellulose membranes using the Thermo Scientific™ Pierce™ G2 Fast Blotter. Following electroblotting, the membranes were blocked using 5% nonfat milk in TBST for 1 hour at room temperature. Membranes were then gently washed with TBST and incubated with Anti-IL-1 beta primary antibody (ab254360, Abcam) at 4°C overnight with continuous rotation. Additional washes, 10 min. 3 times in TBST followed. Before the membranes were probed with Goat anti-Rabbit IgG (H+L) Secondary Antibody, HRP (31460, Thermo Fisher Scientific, RRID: AB_228341) diluted at 1:5000 at room temperature for 1 hour. The membrane was then washed in TBST and immersed into a chemiluminescent HRP substrate solution (Bio-Rad) and imaged using a ChemiDoc MP System (Bio-Rad).

### Immunofluorescent staining of cells

Cells were washed with PBS 3 times, then fixed with 4% PFA for 15 min. Cells were then washed with PBS and permeabilized with 0.1% Triton X-100 in PBS for 20 min. After that, cells were washed with PBS and blocked with 5% normal goat serum (16210072, Gibco) for 1 hour at room temperature. Cells were then incubated with Piezo1 primary antibody (15939-1-AP, Proteintech, RRID:AB_2231460) diluted 1:200, at 4℃ overnight and washed with PBS 3 times. After that, cells were incubated with Goat anti-rabbit Adsorbed Secondary Antibody, Alexa Fluor 488 (A-11008, Thermo Fisher Scientific) at room temperature for 1 hour.

### Actin polymerization staining

Green fluorescent phalloidin conjugate (ab112125, Abcam) was used to stain F-actin according to the manufacturer’s recommendation. Cells were then washed with PBS 3 times and sealed with mounting medium containing DAPI. Data acquisition was performed using a Leica TCS SPE Confocal Microscope.

### Quantification and statistical analysis

Statistical tests were performed using GraphPad Prism 8, which was also used to prepare the graphs. Bars charts represent the mean of the indicated number of experiments or subjects, unless otherwise indicated. Details of the statistical analyses performed for each figure are provided in the figure legends. A p value of < 0.05 or below was considered statistically significant for all experiments.

## Resource availability

### Lead contact

Further information and request for reagents and resources should be addressed to, and will be fulfilled by, the Lead Contact, Lei Sun.

### Materials availability

This study did not generate new unique reagents.

### Data and code availability

All data reported in this paper will be shared by the lead contact upon request, unless it is protected by law.

This paper does not report original code.

Any additional information required to reanalyze the data reported in this paper is available from the lead contact upon request.

## Acknowledgements

This work was supported by the Hong Kong Research Grants Council General Research Fund (15104520, 15102417 and 15326416), Hong Kong Innovation Technology Fund (MRP/018/18X and MHP/014/19), Key-Area Research and Development Program of Guangdong Province (2018B030331001), internal funding from the Hong Kong Polytechnic University (1-ZE1K and 1-ZVW8), and Research Institute of Smart Ageing (1-CD76). The authors would like to thank the facility and technical support from University Research Facility in Life Sciences (ULS) and University Research Facility in Behavioral and Systems Neuroscience (UBSN) of The Hong Kong Polytechnic University. We thank to Dr. Michael Yuen from ULS for instructing us in primary microglia isolation. We thank to Mr. Kai Tang for instructing us in PA gel fabrication.

## Author contributions

T.Z. and L.S. designed research. T.Z., S.K., Y.W., H.C., JJ.Z., C.P.C., XY.Z., T.L. performed research, T.Z., S.K., JH.G., K.F.W., XH.H., MY.Y. analyzed data. T.Z., S.K., and L.S. wrote the paper.

## Competing interests

The authors declare that they have no competing interest.

## Supplementary Figures

**Fig S1.**
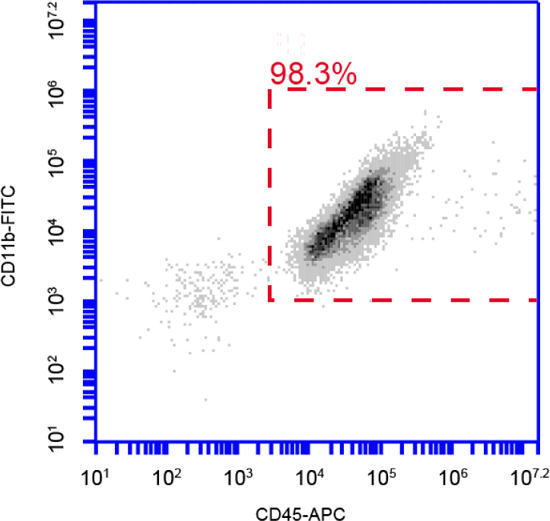
Purity of isolated primary microglia. The purity of isolated primary microglia were determined by flow cytometry with Cd11b-FITC and Cd45-APC staining analyzed by BD Accuri C6 software.

**Fig S2.**
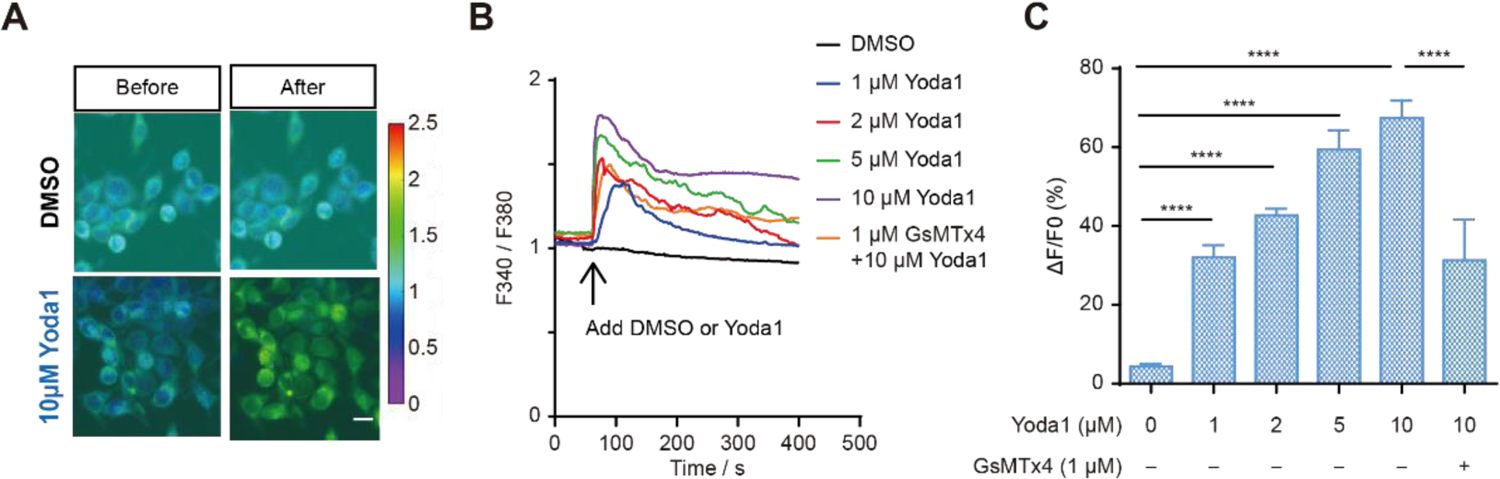
Piezo1 is expressed in the microglial cell line BV2. **A.** Representative live-cell imaging of BV2 loaded with Fura-2 calcium indicator at rest state (left panel) and at the time of maximum fluorescence (right panel). Scale bar= 20μm. **B.** Representative time course of calcium fluorescence in BV2 cells treated with different concentrations of Yoda1 and if pretreated with GsMTx4. **C.** Statistical analysis of ΔF/F0 (%) in BV2 cells treated with different concentrations of Yoda1 and if pretreated with GsMTx4. Bars represent mean ± SEM, N=3 independent experiments; 30–40 cells were analyzed for each experiment. Two-tailed Mann–Whitney U test. ****p < 0.0001.

**Fig S3.**
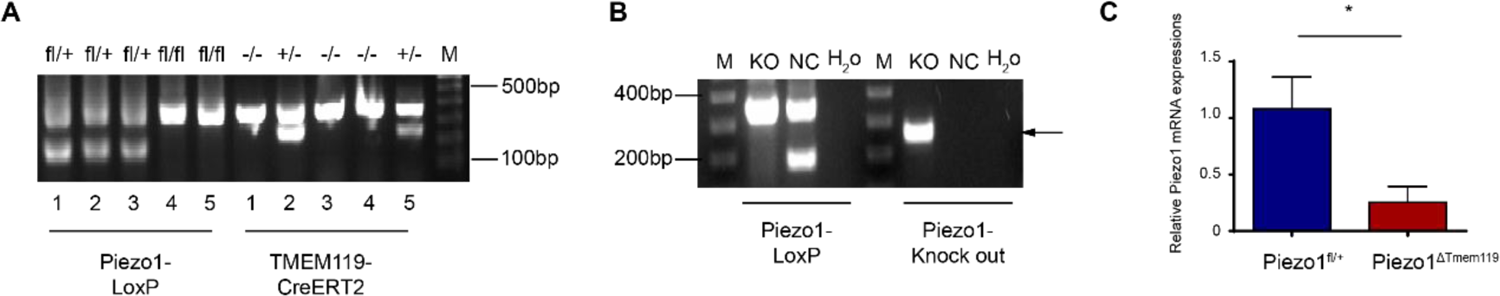
Validation of Tmem119-Piezo1 cKO mice. **A.** Genotyping results of the transgenic mice. 1: Mouse1, Piezo1 fl/+, Cre-/-. 2: Mouse2, Piezo1 fl/+, Cre+/-. 3: Mouse3, Piezo1 fl/+, Cre-/-. 4: Mouse4, Piezo1 fl/fl, Cre-/-. 5: Mouse5, Piezo1 fl/fl, Cre+/-. M represents for DNA ladder. **B.** Piezo1 knock-out validation band. Arrow indicates the Piezo1 knock-out band. KO represents for Piezo1^ΔTmem119^ mouse and NC represents for Piezo1^fl/+^ mouse. H_2_O is adopted as negative control. M represents the DNA ladder. **C.** Piezo1 mRNA expressions (n= 3 in Piezo1^fl/+^ group, n= 4 in Piezo1^ΔTmem119^ group) of brain tissues isolated from Control and Tmem119-Piezo1 cKO mice). Transcript levels were calculated using the 2−ΔΔCT method using β-actin as a reference gene and were normalized to the average expression from control samples. Bars represent mean ± SD. Two-tailed unpaired t-test. *p < 0.05.

**Fig S4.**
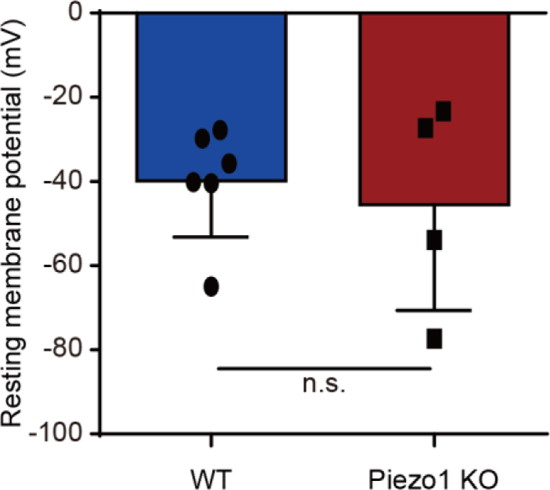
Resting membrane potential of BV2 WT or BV2-Piezo1 KO cells. N=6 cells for BV2 WT group and N=4 cells for BV2-Piezo1 KO group. Bars represent mean ± SEM. Two-tailed unpaired t-test. n.s., no significance.

**Fig S5.**
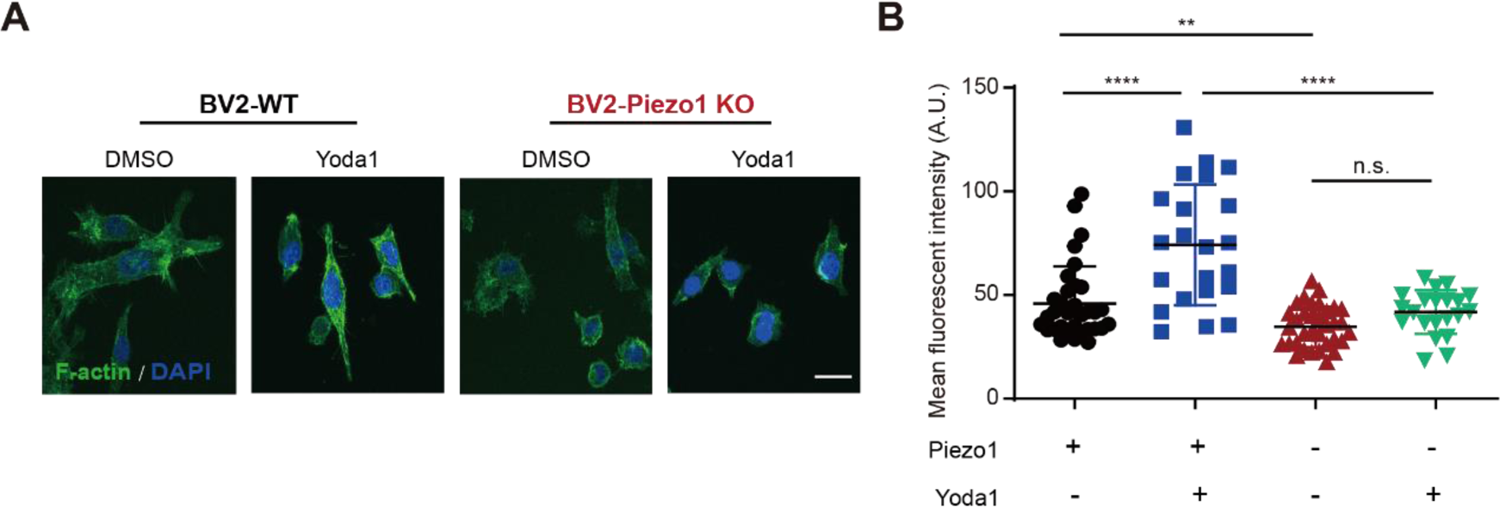
Piezo1 mediates actin polymerization of microglia. **A.** Representative confocal images of F-actin staining (green) and DAPI (blue) under Yoda1 (10μM) stimulation in BV2-WT and BV2-Piezo1 KO cells. Scale bar= 20μm. Quantification of F-actin mean fluorescent intensity (MFI) of BV2-WT and BV2-Piezo1 KO cells stimulated by Yoda1 (10μ M). Repeated for 3 times. 20-35 cells were analyzed for each group. Bars represent mean ± SD. Two-tailed unpaired t-test. n.s., no significance; *p < 0.05, **p < 0.01, ***p < 0.001, ****p < 0.0001.

## Notes

### Competing Interest Statement

The authors have declared no competing interest.

